# Revealing Spatiotemporal Circuit Information of Olfactory Bulb in Large-scale Neural Recordings

**DOI:** 10.1101/2021.03.05.434081

**Authors:** Xin Hu, Shahrukh Khanzada, Diana Klütsch, Federico Calegari, Hayder Amin

## Abstract

Large-scale multi-site biosensors are essential to probe the olfactory bulb (OB) circuitry for understanding the spatiotemporal dynamics of simultaneous discharge patterns. Current ex-vivo electrophysiological techniques are limited to recording a small set of neurons and cannot provide an inadequate resolution, which hinders revealing the fast dynamic underlying the information coding mechanisms in the OB circuit. Here, we demonstrate a novel biohybrid OB-CMOS platform to decipher the cross-scale dynamics of OB electrogenesis and quantify the distinct neuronal coding properties. The approach with 4096-microelectrodes offers a non-invasive, label-free, bioelectrical imaging to decode simultaneous firing patterns from thousands of connected neuronal ensembles in acute OB slices. The platform can measure spontaneous and drug-induced extracellular field potential activity. We employ our OB-CMOS recordings to perform multidimensional analysis to instantiate specific neurophysiological metrics underlying the olfactory spatiotemporal coding that emerged from the OB interconnected layers. Our results delineate the computational implications of large-scale activity patterns in functional olfactory processing. The high-content characterization of the olfactory circuit could benefit better functional interrogations of the olfactory spatiotemporal coding, connectivity mapping, and, further, the designing of reliable and advanced olfactory cell-based biosensors for diagnostic biomarkers and drug discovery.

## INTRODUCTION

The olfactory bulb (OB) is a vital chemosensory structure of the information coding mechanisms^1^. It allows vertebrates to process and discriminate vast complex odorants and distinguish them with high selectivity and sensitivity^2,3^. The OB’s neuronal processes are distributed intricately, composed of an abundance of dendrodendritic interactions and a stratified structure in five connected layers^4,5^. The OB also renders a distinctive form of morpho-functional neuronal plasticity conferred by a constant supply of new neurons (i.e., adult neurogenesis), allowing profound remodeling of the bulbar circuit in response to experience and challenges^6^.

This organization represents a high degree of plasticity and is the first relay station of olfactory perception with downstream information processing from the primary sensory epithelium to high-order neurons in subcortical and cortical areas for odor identification and interpretation^7–9^.

This high dimensionality of olfactory processing and odor coding properties inspired the development of biomimetic olfactory sensors, i.e., electronic noses^10–12^. They promise potential advances in medical diagnosis, food quality, environmental, and military applications^13^. However, a fundamental inherent shortfall in these electronic nose systems is the lack of realistic dynamics at the physiological cellular and network levels, thus hindering this technology’s exploitation for further challenging applications^14^. Hence, considerable achievements in genetic, biotechnology, and bioengineering have been put forth to enable the implementation of olfactory cell-based biosensors (i.e., bioelectronic noses)^14,15^ to circumvent the limitations of conventional electronic noses and to promote specific measurement of target odorants^16^. In this context, many studies have been reported to employ a range of different olfactory cell-based biosensors^16–18^. Notably, a roadmap for advancing this technology has focused on deciphering the inherent plasticity mechanisms of olfactory information encoded in the orchestrated spatiotemporal activity patterns to refine and stabilize the OB sensory information^9^. These patterns are rhythmic neuronal synchronizations that encompass time-varying spatial distributions of discrete spike trains superimposed and complemented with a slow oscillation < ~300 Hz of the local field potentials (LFPs) that reflect subthreshold integrative processes^19–21^.

On the other hand, the mammalian olfactory system generates rhythmic oscillations of LFPs that exhibit a functional role in local odor processing and odor discrimination (i.e., gamma frequency band; 40-100 Hz)^22^, odor learning and experience (i.e., beta band; 15-30 Hz)^23^, and respiratory oscillations overlapping with the (theta band; 2-12 Hz)^24^. Thus, investigating the electrophysiological properties is crucial to further understand the local OB functional circuit connections and behavior and how they shape sensory processing. In view of these demands, many previous studies have used cell or slice-based biosensors to report experimental paradigms and classical recording techniques for different olfactory signal biosensing such as microelectrode arrays (MEAs)^25–27^, semiconductor field-effect transistors^28,29^, patch-clamp recordings^30^, light addressable potentiometric sensors^31^, quartz crystal microbalance^32^, electrochemical impedance spectroscopy^33^, and surface Plasmon resonance^34^.

Also, these efforts are aided by insights from computational models^35^ and neuromorphic computing^36^. Despite MEAs shortcomings (i.e., low spatial resolution due to low numbers of recording electrodes), this method enables non-invasive, multi-site, long-term, and label-free probing of neuronal firing activity without disruption of cellular integrity. Recently, the manufacturing of low-cost semiconductor techniques based on complementary metal-oxide-semiconductor (CMOS) technology has been introduced into the design of biosensors^37^ and the implementation of a high-density active-pixel sensor (APS) MEAs^38,39^. Thereafter, APS CMOS-MEAs have been exploited in a wide range of neuroscience and bioengineering applications^40–42^. Their unique inherent features with on-chip addressing and multiplexing and high signal-to-noise ratio (SNR) allow the measurement of neural signals at high spatiotemporal resolution with simultaneous recordings from several thousands of electrodes both in vitro^38,39^ and in vivo^43^. In turn, these features enabled large-scale recordings and network information analysis from the interaction of various neuronal components to facilitate understanding the mechanisms that give rise to behavior-dependent of cell assembly patterns^44^. However, spatiotemporal circuit interactions underlying OB coding mechanisms have remained mostly elusive, hindering the development of advanced olfactory-cell-based sensors. Although existing olfactory-MEA biosensors have endowed valuable results, a clear understanding of the interareal coordination of bulbar circuitry mechanisms has lagged behind.

In this study, we report a novel Biohybrid Olfactory Neural Circuit on a CMOS-chip (BIONICS) capable of recording massively simultaneous neuronal firing patterns in acute mouse OB slices. The platform integrates large and dense electrode arrays (i.e., 4096-microelectrodes) and on-chip signal processing implemented in a CMOS-chip. The combination of the OB circuit and the high-density electrode configuration allows the recording of real-time spontaneous and pharmacologically-evoked activation of the OB neural circuit from thousands of connected neuronal ensembles within the interareal layers of the OB operated in ex vivo. We further exploit our OB-CMOS-chip recordings with fundamental basic and advance analyses to instantiate specific neurophysiological metrics underlying the olfactory spatiotemporal information that emerged from the OB interconnected layers. Therefore, we characterize the rhythmic oscillations of the LFPs, oscillatory frequency bands, and coherence magnitude for single electrode activity as well as between OB layers. We also identify the event initiation onsets and quantify the dynamic profiles of signal propagation patterns. The rich information ingrained in our unique datasets also allows us to compute the spatial neural sources that contribute to the LFPs by using the kernel current source density (kCSD) method, functional connectivity (FC) maps, and topological interaction of thousands of active OB neuronal ensembles that delineate the construction of functional OB circuitry. Finally, we validated our platform to unveil a large-scale functional remodeling in the OB circuitry due to ongoing neurogenesis using a well-established and reported mouse model of enhanced neural stem cell expansion^45–47^.

To our knowledge, this is the first report of a high-density CMOS-neurochip that allows revealing the spatiotemporal functional processing in the interconnected OB circuit. Deciphering the mechanism of large-scale OB electrogenesis will enhance our understanding of the neuronal ensembles’ functional organization. It will also provide an effective tool to investigate information processing and spatiotemporal coding in the OB circuit. Together, our results offer vital evidence for the future development of biomimetic olfaction detection biosensors for various applications.

## MATERIALS AND METHODS

### Animals and OB acute slices preparation

All experiments were performed on 8 weeks C57BL/6j (Charles River Laboratories, Germany) and triple transgenic 4D− and 4D+ mice, previously described^45^. All work and animal procedures were performed in accordance with the applicable European and national regulations (Tierschutzgesetz) and were approved by the local authority (Landesdirektion Sachsen; 25-5131/476/14 and DD24-9168.11-1/2011-11, TVV13/2016, and HD35-9185.81/G-61/15). Mice were anesthetized with isoflurane before decapitation. The brain and OB were carefully removed from the skull and placed in a chilled cutting sucrose solution before slicing. The Brain and OB were fixed on the cutting plate, and horizontal slices (300 μm thick) were prepared using Leica Vibratome VT1200S (Leica Microsystems, Germany). Slices were cut at 0-2°C in aCSF solution saturated with 95% O_2_ and 5% CO_2_ (pH = 7.2-7.4) of a high sucrose solution containing in mM: 250 Sucrose, 10 Glucose, 1.25 NaH_2_PO_4_, 24 NaHCO_3_, 2.5 KCl, 0.5 Ascorbic acid, 4 MgCl_2_, 1.2 MgSO_4_, 0.5 CaCl_2_. Next, OB slices were incubated for 45 min at 34°C and then allowed to recover for at least 1 hour at room temperature before used for recordings with CMOS-chips, in a recording aCSF solution containing in mM: 127 NaCl, 2.5 KCl, 1.25 NaH_2_PO_4_, 24 NaHCO_3_, 25 Glucose, 1.2 MgSO_4_, 2.5 CaCl_2_, and the solution was aerated with 95% O_2_ and 5% CO_2_.

### Biohybrid OB-CMOS biosensor recordings

We performed all extracellular recordings using CMOS-biosensors and an acquisition system (3Brain AG, Switzerland) customized to our BIONICS setup. CMOS-chips integrate 4096 recording electrodes with 42 μm pitch-size to compose an active sensing area of ~7 mm^2^, ideal for recording from the entire ~4 mm^2^ OB tissue. The on-chip amplification circuit allows for 0.1-5kHz bandpass filtering conferred by a global gain of 60 dB sufficient to record slow and fast oscillations. Modular Stereo microscope Leica MZ10F (Leica Microsystems, Germany) was designed and incorporated into the setup to capture OB slices’ light-imaging simultaneously with the whole-circuit firing pattern recordings. All CMOS-chips were coated with 0.1 mg/ml PDLO (Sigma-Aldrich, Germany) and incubated for 30 min at 37°C before recordings. Then, slices were moved onto chips, and the coupling between OB slices and CMOS-electrodes was enhanced with a custom-made platinum harp placed above the tissue. Slices were perfused continuously with oxygenated recording solution at 4 ml/min. We collected 10 min of spontaneous and pharmacological-induced extracellular recordings at 14 kHz/electrode sampling frequency and stored them for offline analysis. We used Brainwave software (3Brain AG, Switzerland) for data recording and employed a hard threshold algorithm to perform LFP detection with a 10 ms refractory period and a 500 ms maximum event duration.

### Drug treatment

We prepared fresh solutions of all drugs for each experiment by dissolving the drug in the recording solution. The final working concentrations used for all experiments were 100 μM, 30 μM, and 10 μM of 4AP, BiC, and MK-801 (Sigma-Aldrich, Germany), respectively.

### Immunohistochemistry and fluorescence imaging

Brains with OB were perfused and post-fixed overnight in 4% PFA at 4°C. Immunohistochemistry was performed as described^45^. The images were acquired with an automated Zeiss ApoTome confocal microscope (LSM 780, Carl Zeiss, Germany).

### Data Analysis

All basic and advanced algorithms used in this study were developed as custom-written Python scripts.

### Mean activity basic analysis

We selected four parameters to describe the mean activity features of large-scale spatiotemporal LFP oscillations, including amplitude, energy, frequency, and duration. The signal amplitude analysis was obtained by full-wave rectification and low-pass filtering (cutoff frequency 100 Hz). The energy is defined as the area under the squared magnitude of the LFP rates for a specific time window.

### Lognormal distribution

The LFP firing patterns of the OB neuronal ensembles show a wide degree of participation in the circuit activity and followed a skewed lognormal distribution. Thus, we computed the probability density function for the lognormal distribution:

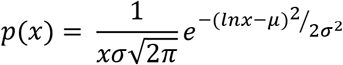

Where *μ* is the mean, and *σ* is the standard deviation.

### Gini coefficient and CV2

To quantify the inequality of participation of individual neurons in the OB circuit, we employed the Gini index as a commonly used measure of inequality and sparsity^48^ and computed as the ratio of the areas on the Lorenz curve diagram:

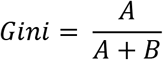

Where *A* is the area above the Lorenz curve, and *B* is the area below for the cumulative LFP firing rates.

To assess the local-trial variability of the firing patterns, we used the coefficient of variation (CV2)^49^, which is defined as the function of pairs of adjacent inter-LFP-event intervals *IEI_i_* and *IEI*_*i*+1_ divided by the average of the event intervals.

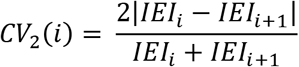

### Time-frequency and PSD analyses

We first constructed the frequency-time dynamics in pseudo-color spectrograms for a selected 10 sec time window using a filtered LFPs (1-100 Hz). Next, we computed the Periodograms to identify dominant frequencies in the oscillatory activity in a given time-series. The spectra were calculated using Welch’s method^50^ by calculating the Fast Fourier Transform of the recorded LFPs to estimate the power spectral density (PSD).

### Waveform classification

To characterize and allocate the nonsinusoidal shapes of the recorded LFP waveforms to their OB layers, we developed a procedure consisting of four steps. First, (waveform extraction) - we detected active firing electrodes clustered based on their structural relation to the OB layers. We extracted information from hundreds of waveform segments that occurred at least in three consecutive LFP events. Second, (Pre-processing) – all detected and extracted waveforms were denoised by hard threshold algorithm and filtered by Low pass filter (1-100 Hz). Third, (Parameter Identification) – we defined in our extracted waveforms four features, including amplitude (waveform height feature), period, rise-decay symmetry (slope features), and area (area feature). Fourth, (classification and clustering), we reduced the dimensionality of the computed waveform features using the Principal Component Analysis (PCA) and K-means clustering algorithms^51^ to identify best-classified waveform shapes into five clusters associated with the GL, GCL, PL, ONL, and OCx layers.

### Spatial coherence

To measure the predictable relationship between firing LFP rates at different spatial locations of the OB circuit, we calculated the 2D spatial coherence maps. We calculated the correlation between pairs of firing electrodes of the CMOS-chip (64 × 64 electrodes) arranged in rows (x) and columns (y). The function is a dimensionless quantity and ranges between (0,1) and defined as^52^:

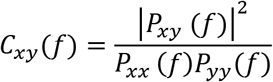

Where *P_xx_*(*f*) and *P_yy_*(*f*) are the autospectral density of the firing x and y, respectively, and *P_xy_*(*f*) is the cross-spectrum density of the firing x and y estimated using Welch’s method. A Spatial coherence near one indicates that the active electrodes encode all information of the firing rate of that spectral frequency component. In contrast, a value close to zero indicates no information of the firing is present.

### LFP propagation dynamics

To quantify the propagation magnitude of the spatiotemporal LFP Events in all interconnected OB layers, we employed the center-of-activity trajectories (CATs) analysis^53,54^. We used the voltage values embedded in the LFP frame activities within 5 ms moving time bins to collect the CAT magnitudes in each firing event. The value of the CA at time *t* is a two-dimensional vector defined as:

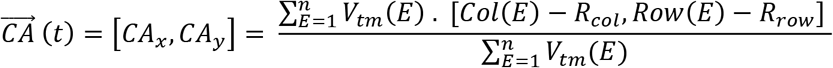

Where *V_tm_*(*E*) represents the LFP firing rate corresponding to the active electrodes *E* within a time window (*tm*). Also, *Col*(*E*) and *Row*(*E*) are the column and row numbers of the associated *E.* Then, *R_col_* and *R_row_* are the coordinates of the physical center of 64 × 64 electrodes. *n* is the total number of active electrodes. Then, the CA trajectory from *t*_0_ to *t*_1_ with a time step Δ*t* is defined as:

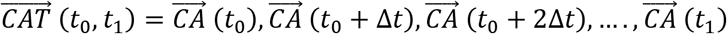

### Velocity of conduction

To track the putative displacement of 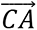 of LFP events in the entire interconnected OB circuit, the velocity of conduction was calculated as the average of instantaneous vector quantity displacements calculated for the time of a propagating LFP event and expressed as:

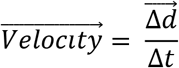

Where *Δd* is the change in displacement and *Δt* is the change in time.

### Functional connectivity and causal links

To infer the large-scale statistical dependent connectivity in a multilayered OB network, we first calculated the cross-covariance between pairs of active electrodes in the 64 × 64 array using the Pearson correlation coefficient (PCC)^55^. The correlation coefficient between electrodes was then sorted based on the OB layers and presented in a symmetric matrix. The PCC is defined as:

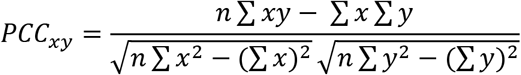

Where *x, y* denoted the values of pair electrodes time series for *n* number of the active array.

Then, we performed the Multivariate Granger causality by fitting a vector autoregressive model to the time series to quantify the influence of one time series on another^56^. To measure the directional information flow within the correlated links in the network, we performed the Directed Transfer Function (DTF)^57^.

### Current source density

To estimate the current sources generating the extracellular low-frequency potentials recorded in all OB layers, we used the Kernel Current Source Density (kCSD) Analysis^58^, described in the kCSD-python package and available on GitHub (https://github.com/Neuroinflab/kCSD-python).

### Network efficiency and wiring cost

To quantitatively measure a precise information flow and exchange of parallel integrated processing in the interconnected OB network, we used the measure of global, local efficiencies^59^, and connection cost. These are dimensionless measures ranging from 0 to 1. The global efficiency can be defined as the inverse of the average of shortest path lengths between all nodes in the network and calculated as:

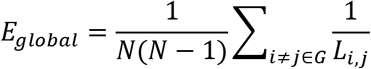

Where *N* is the number of all nodes in the network, *L_i,j_* is the average path length between all nodes in the network.

Further, based on local clusters, the local efficiency is the average efficiency of clusters (subgraph) for a node with the neighbors, and calculated as:

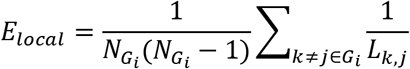

Where *N_G_i__*. is the number of nodes in the local subgraphs (clusters) *G_i_* of the network, *L_k,j_* is the path length between pair nodes in the cluster.

### Wiring cost

The network cost was estimated based on the minimum spanning tree (MST) method using the Kruskal algorithm. This approach is calculating the shortest path length connecting all network nodes at the cheapest cost.

### Statistical analysis

All statistical analyses were performed with Originlab 2020. All data in this work were expressed as the mean ± standard error of the mean (SEM). All box charts are determined by the 25^th^-75^th^ percentiles, and the whiskers by the 5^th^-95^th^ percentiles, and lengths within the Interquartile range (1.5 IQR). Also, the lines depict the median and the squares for the mean values. Differences between groups were examined for statistical significance, where appropriate, using one-way analysis of variance (ANOVA) or two-way ANOVA followed by Tukey’s posthoc testing. *P* < 0.05 was considered significant.

## RESULTS AND DISCUSSION

In what follows, we first developed and implemented a hybrid biosensor platform by integrating the OB circuit into a CMOS-MEA chip, which allowed us to record LFPs and spikes from the OB multilayers. Next, we exploited the multidimensional data focusing on the LFPs to infer and provide new insights about the subthreshold integrative processes in high spatiotemporal resolution expressed by extracting unique dynamic features of the oscillatory activity and functional mapping across the OB circuitry. (i.e., frequency spectrum, coherence, cross-correlation, causal connectivity, and assessing neurogenic remodeling effect).

### Implementation of the large-scale BIONICS platform

To quantitatively probe the large-scale firing patterns and characterize the olfactory spatiotemporal coding of the whole-multilayered, we engineered a biohybrid OB-CMOS biosensor platform (BIONICS) (**Figure 1**). It combines isolated whole-OB acute slices and a CMOS-chip. The OB-CMOS-chip continuously perfused with an optimized aCSF to maintain high viability and stable, functional responses for several hours (*n=24 slices from 8 adult mice at 8 weeks old*). Also, we employed optical imaging into the bioelectrical OB-CMOS-based setup allowing the light-imaging acquisition of OB slices concurrently with the whole-circuit firing pattern recordings **(Figure 1a**). The CMOS-chip integrates a 64 × 64 microelectrodes pixel array to allow the simultaneous recordings of sub-millisecond extracellular firing information from the entire OB circuitry (**Figure 1b**). The active sensing area of the CMOS-chip is 2.69 × 2.69 mm (i.e., 4096 microelectrodes with 42 μm pitch)^39^, which is compatible with the area of a whole mouse OB slice (i.e., 2 × 2 mm). Our OB-CMOS biosensor readout captured the real-time circuitry acquisition rendered by a single electrode (i.e., pixel-like) sensor from sequential bioimaging frames encoding olfactory functional information at high spatiotemporal resolution. The voltage values of each sensing pixel were mapped on large-scale OB layers with a pseudo-color scaling. The final construction illustrated real-time bioimaging video frames of the entire OB functional circuit (**Figure. 1b**, and **Supplementary Movie 1**). The CMOS-chip surface was functionalized using a specific adhesion-promoting molecule – poly-DL-ornithine (PDLO)^60^ that enhanced the cell-electrode interface coupling and improved SNR by increasing the seal resistance (*R_seal_*) between the tissue and the electrode (**Figure 1c**).

**Figure 1.**
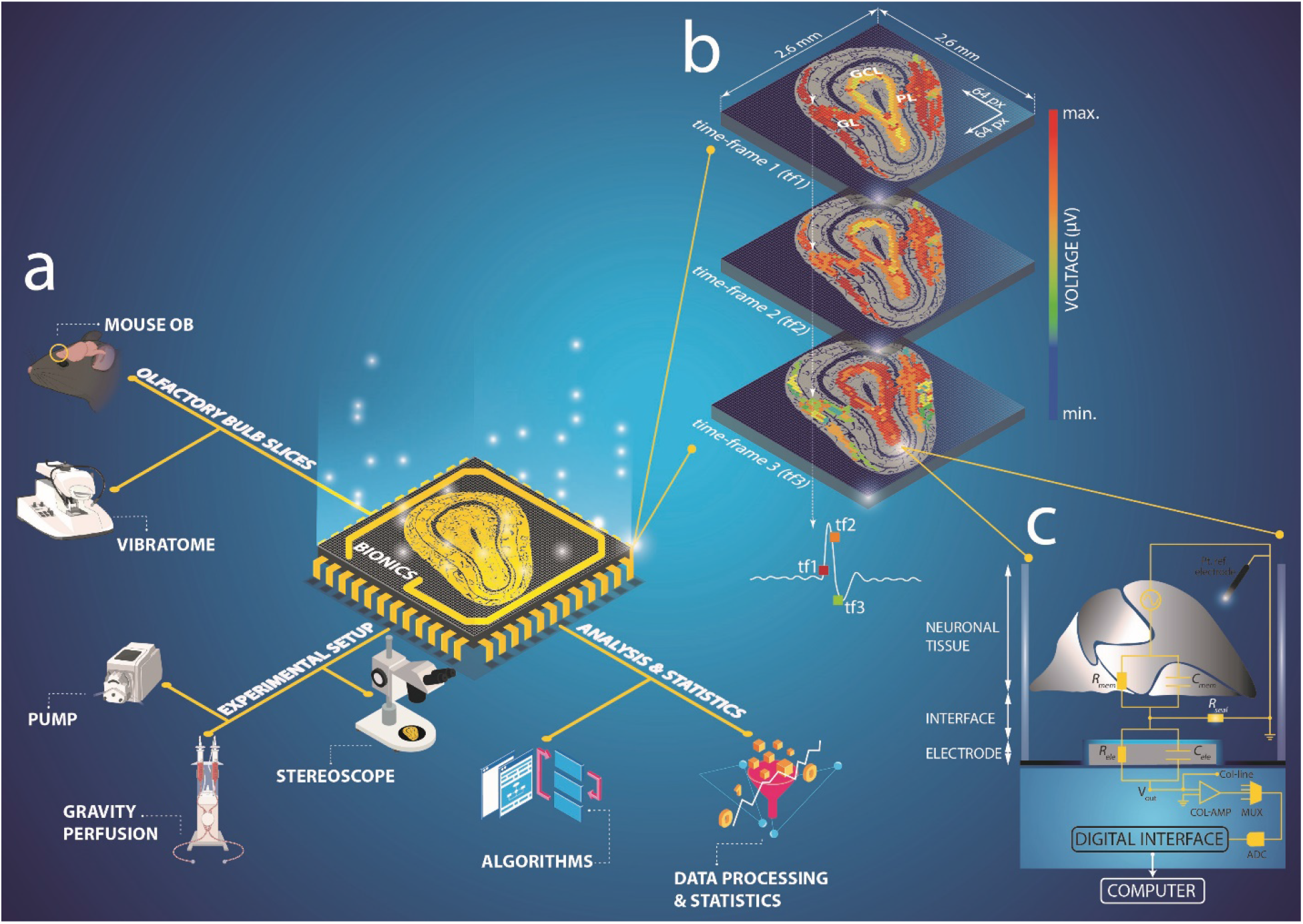
BIONICS setup and implementation. **a)** Graphical isometric imaging setup featuring the biohybrid OB-CMOS-chip with 64×64 electrode array and integrating the experimental and analysis components. **b)** Pixel-multi-frame real-time representation of the encoded OB network-wide activity enabling pseudo-color reconstruction of the firing information from the sequential obtained frames. **c)** Cell model of tissue-electrode CMOS interface showing the pixel circuit configuration, the individual functional elements, and the communication interfaces. Surface functionalization with PDLO enhanced the slice-electrode coupling interface and increased the SNR (i.e., increased the *R_seal_*).

### Features of spatiotemporal LFP firing patterns and activity-dependent distributions in multilayered OB circuits

The neuronal processes in the OB instantiate in five connected layers, i.e., olfactory nerve layer (ONL), glomerular layer (GL), external plexiform layer (EPL), mitral cell layer (MCL), and granule cell layer (GCL)^4,5^. In our study, we consider the EPL and the MCL as the projection layer (PL). In a canonical biological olfactory circuit, the olfactory sensory neurons (OSNs) project and converge their axons into the same glomeruli in the GL. Glomeruli represent the inhibitory functional units for information coding that shape the features of odorant responses^61^. OSNs form excitatory synapses with mitral cells in the projection layer that, in turn, project their axons to the olfactory cortex (OCx)^62^.

To address the overarching benefit of the OB-CMOS-biosensor to reveal different neuronal processes by their intrinsically generated firing patterns or the facilitation of synaptic transmission, we recorded the spontaneous and pharmacologically-induced large-scale responses in OB circuitry. We used three biochemical benchmark compounds including, 4-aminopyridine (4-AP) - potassium channel blocker, Bicuculline (BiC) - GABA_A_ antagonist, and dizocilpine (MK-801) - N-Methyl-D-aspartate (NMDA) receptor antagonist. We computed several first-order statistical parameters (LFP energy, oscillation frequency, amplitude, and duration). Spontaneous oscillation and spiking activity were detected in all OB layers except the ONL. We found substantial responses of the OB circuit assessed by the statistical parameters shown in (**Figure 2a, *right***). We also illustrated the spatial organization of those large-scale metrics by overlaying the computed mean of the functional, statistical metrics obtained from the LFP recordings on the OB circuits’ optical images (**Figure. 2a *left***).

**Figure 2.**
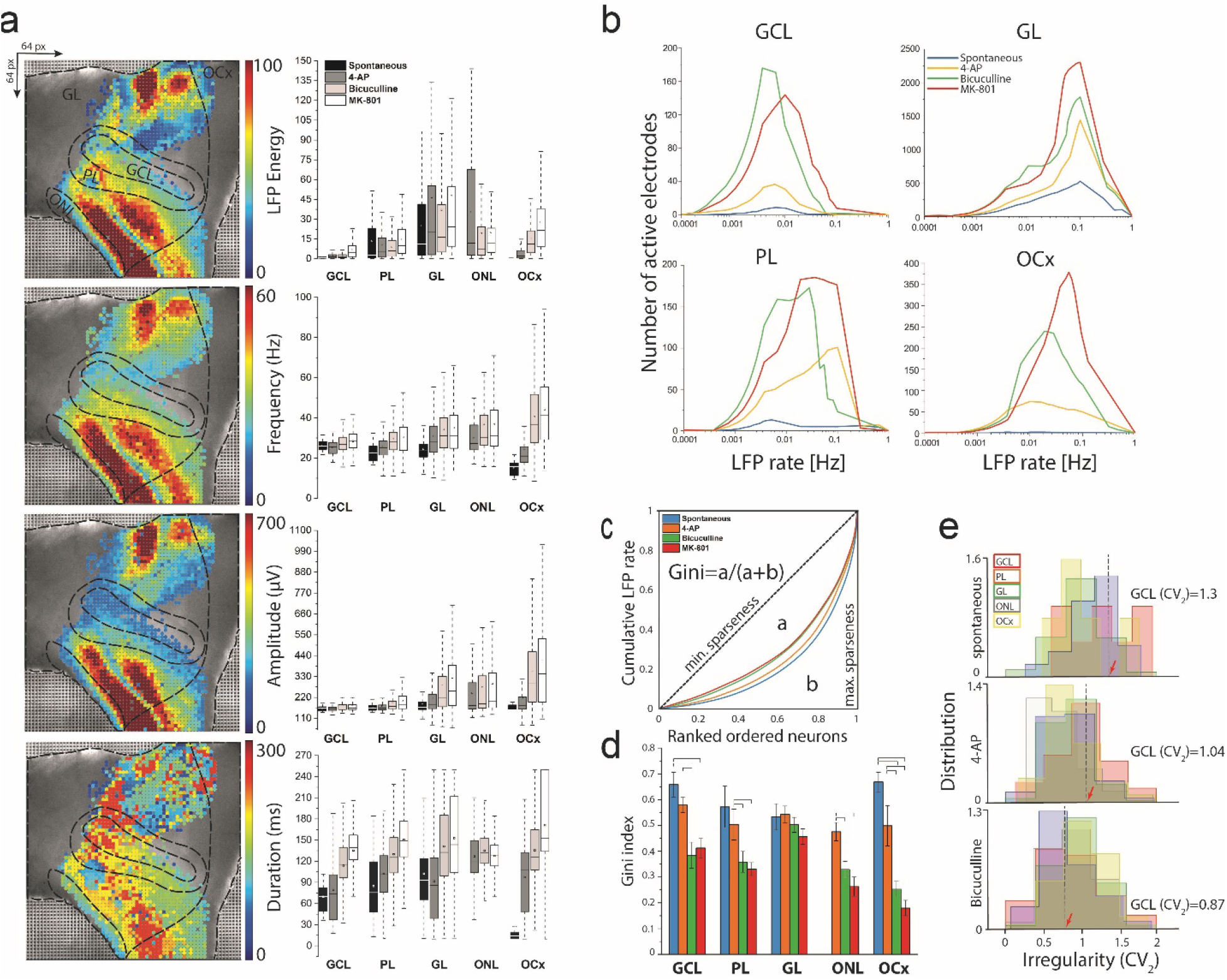
OB network-wide extracellular activity parameters and distribution. **a)** Topographical representations and statistical characteristics of the large-scale firing patterns recorded simultaneously from the entire OB slices (i.e., 4AP induced activity example). Pseudo-color values of the energy, frequency, amplitude, and LFP activity duration superimposed to the OB structural images. LFP energy is significantly higher in GL than other layers, and frequency denotes a similar frequency oscillation range in all layers. The amplitude is also significantly higher in GL and increased with drug treatment; in contrast, the LFP events’ duration is decreased with consecutive drug treatments. **b)** LFP firing events display a lognormal-like distribution in interconnected OB layers allowing to elucidate the network dynamics. **c)** Lorenze curve and Gini coefficient analysis indicate the inequal neuronal participation and sparseness. **d)** Gini coefficient quantification for the firing activity in all OB layers (\**p* < 0.05, \*\**p* < 0.01, ANOVA). **e)** CV_2_ values (0 → 2) showing the quantification of the spread of IEIs for the entire firing events. The CV2 is reduced (towards regular pattern) after blocking the GABAergic population with BiC as indicated by moving red arrows to the left side (\*\**p* < 0.01, ANOVA) (*n=24 slices from 8 mice*).

The collective functional representations given in firing patterns of synchronized neuronal populations have been shown to exhibit a skewed distribution of firing rates in other experimental setups^41,63^. Our large-scale OB recordings found similar highly skewed lognormal-like distributions of firing LFPs in GCL, Pl, and OCx layers (**Figure 2b**). Those skewed distributions showed a range of low and high firing patterns indicating a network-wide firing fluctuation and a diverse repertoire of participating neurons to the circuitry firing information. In other words, when the OB circuit activity was enhanced by unbalancing the inhibition or excitation drive (i.e., BiC and MK-801 treatments), the firing distribution showed lognormal-like distributions in the GCL, PL, and OCx, but a symmetric distribution in the GL, indicating a mixture of circuit firing regimes of fluctuation-driven and mean-driven, respectively^64^. Remarkably, this mixture of activity regimes within interconnected OB layers indicates the significance of featured spatiotemporal patterns in maintaining the sensitivity and stability balance of the OB circuitry.

To parametrically evaluate the spatial sparsity and diversity of participating neurons (i.e., the efficiency of neuronal representations) rendered by their firing potentials, we used Lorenz statistics to generate the Gini coefficient^48^. The Gini index is between zero and one and reflects the inequality of participation of neurons giving by their firing LFP values; it is higher in the unequal participation of neurons to the firing activity (**Figure 2c**). The Gini index was higher in the spontaneous recorded LFPs in all OB layers (i.e., particularity in GCL, PL, and OCx), which decreased along with the treatment of the compound of 4-AP, BiC, and Mk-801, respectively (**Figure 2d**). This indicated that neuronal selectivity and the sparseness of representation in a given spontaneous activity reflected the strong activation of a relatively small group of cells. This is the first evidence of the sloppiness in spontaneously active OB layers. A small but stable subnetwork of neurons is critical for global stability, allowing considerable plasticity to occur in the remaining majority of cell population^65^.

On the other hand, the evoked-activity groups (i.e., BiC and MK-801) showed sloppiness but not sparseness, which inferred stiff dimensions involving more complicated combinations of other parameters following the circuit modulation. Despite the overall stability we observed in the OB functional representations, changes in the LFP timings were found upon manipulating the balance of inhibition and excitation by pharmacological treatment. To evaluate these changes in all OB layers, we quantified the LFP timings irregularity using the coefficient of variation index (CV2)^49^(see methods). We found a significant decrease in the CV2 (i.e., GCL=1.3, 1.04, and 0.87) in spontaneous, 4-AP, and BiC conditions, respectively (**Figure 2e**). This decrease in CV2 values indicates decreased variability, hence, increased LFP regularity upon compound treatment. Altogether, our results delineate detailed spatiotemporal functional implications of discharge patterns in distinct subnetworks of the OB layers opt for stability and spatial information coding. This could emerge from the intrinsic biophysical properties of the individual ensembles in these circuits and their distinct wiring, as reported in other brain states^63^.

### Characterization of oscillatory-dependent features of LFPs and waveform classification

Next, we sought to identify the functional significance in the frequency and time domains of the LFP oscillatory synchronized waves rendered by the interconnected OB layers. Thus, we employed band-pass filters on the recorded data to set for high-frequency oscillation (HFO), indicating spiking activity and low-frequency-oscillation (LFO), indicating three prominent oscillatory bands in different layers (i.e., theta, beta, and gamma) **(Figure 3a)**. We observed slow and fast oscillations in nearly all layers under spontaneous and pharmacologically-modulated activity (**Figure 3a**, GL and GCL layers examples). Then, we computed the synchronous frequency-time domain dynamics of a selected electrode within the 4096-microelectrode array in pseudo-color spectrograms. This spectrogram exhibited superposed sustained oscillatory events in the GL layer lasting for several seconds with coordinated frequency peaks in the theta-gamma band **(Figure 3b, *left***).

**Figure 3.**
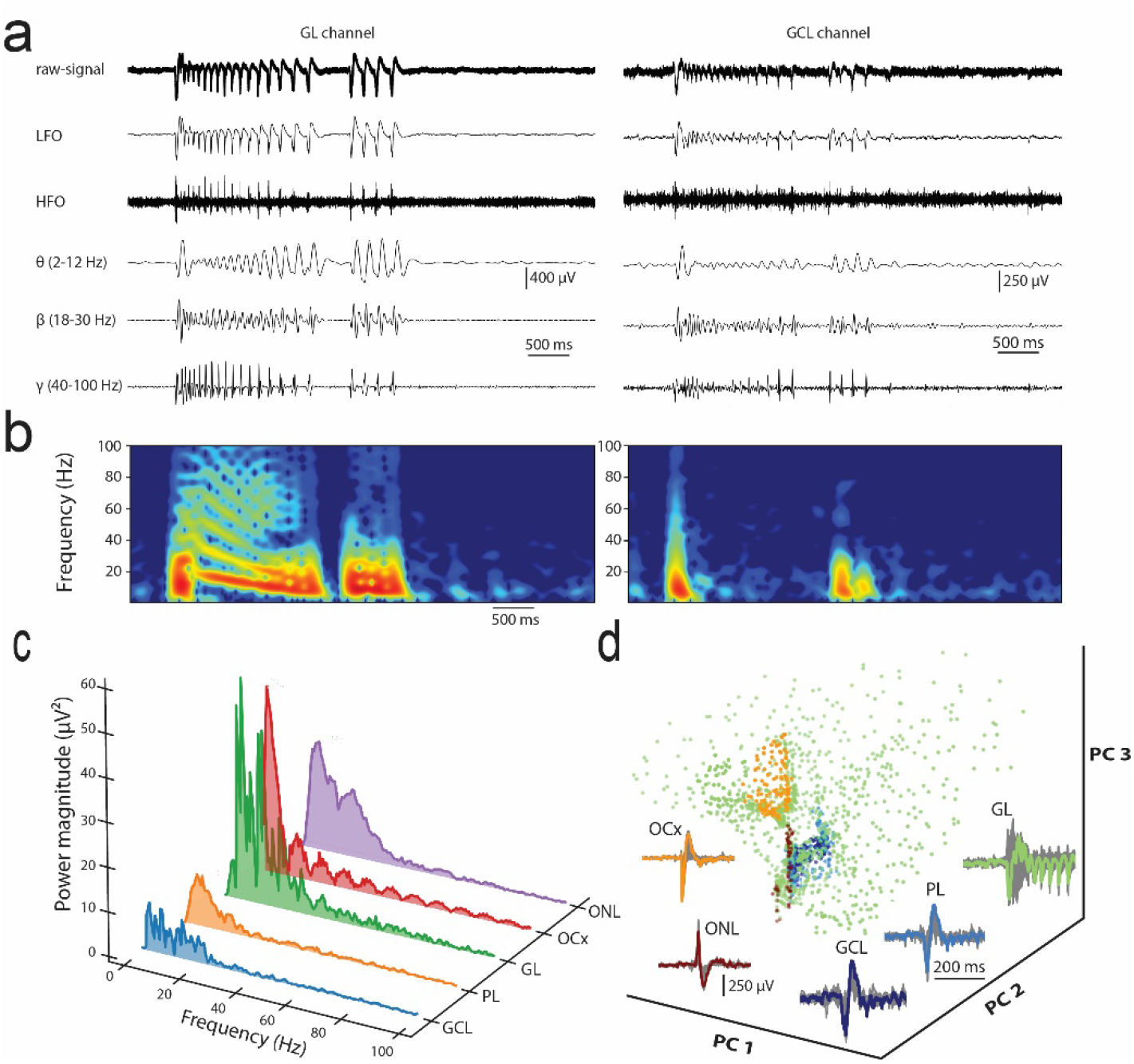
Network-wide oscillation features and waveforms signatures. **a)** Representative event traces from large-scale CMOS-chip recording sites showing low to high ranges of oscillatory frequencies in GL and GCL layers. Exemplary signals from the GL and GCL show wide biosignal signatures of spikes and LFPs with frequency components (θ, β, and γ bands). **b)** Pseudo-color spectrograms showing the Frequency-time dynamics in GL and GCL layers. **c)** Representative power spectral density analysis depicts the LFP signals’ strength regarding their broadband oscillation in OB layers. It displays a high power magnitude of the θ band in the GL. **d)** Classification of multilayered OB oscillatory waveform shapes profiled with the PCA analysis and K-means algorithm based on features extracted from the LFP events. Superimposed colored average waveforms correspond to the OB layers, and grey waveforms showing all extracted extracellular waveforms.

Similarly, **(Figure 3b, *right*)** shows the GCL layer’s spectrogram, where single transient pulses of oscillatory events occurred with a frequency range between 4-60 Hz. Further, we employed the power spectral density (PSD) analysis to simultaneously quantify a specific oscillatory band’s power magnitude in the interconnected OB layers recorded by a full-CMOS array from spontaneous and drug-induced activity phases **(Figure 3c,** and **Supplementary figure 1).** Despite the noticeable fluctuations in the power magnitude upon pharmacological manipulation, we found a prevailing network frequency oscillation in the theta band in all OB layers, predominantly in GCL and GL. These results confirm evidence on the rhythmic theta oscillations coordinated by the glomerular network^66^. Further, the theta-gamma coupled oscillations were also reported to emerge from the individual mitral cells^67^ and also found in hippocampal networks^68^. Our results suggest rhythmogenic subregions of the OB circuitry to drive distinct functional microcircuits that are vital for OB coding.

In addition, investigating the features of oscillatory waveforms can better elucidate the contribution of individual neuronal populations in the spatially connected layers and the olfactory information coding. Therefore, we employed unsupervised clustering of LFP waveforms, which resulted in five distinct waveform classes across all OB layers **(Figure 3d)**. We analyzed hundreds of waveforms in each OB layer made possible by the multidimensional high spatiotemporal resolution CMOS-recordings. The classification process was performed by PCA and clustering with the k-means algorithm (see methods).

### Mapping the spatial firing patterns for global circuit dynamics

Coherent oscillatory activity in the olfactory circuit is rendered by the spatiotemporal LFP patterning across the OB layers. We sought to exploit BIONICS to quantify the spatial mapping of large-scale firing rate encoded in the whole OB interconnected layers. To do so, we computed the spatial coherence as the correlation of the encoded firing rates of interelectrode distance. The spatial extent of LFP rates is derived from each pixel in the 4096-microelectrode array to capture the dynamics of rhythmic components in space and estimate the breadth of activity origin. Thus, higher coherent electrodes have more regulated LFP firing in a specific cluster of the OB region in space, while lower coherent electrodes show more nonstationary and lower firing activity. In all recorded OB slices, coherence increased across evoked-activity conditions (spontaneous vs. 4AP, BiC, and MK-801, respectively) due to increased spatial rate coding under the activation of more cells in the OB circuits **(Figure 4a)**. We also computed and quantified the population average of the spatial coherence across the pharmacological conditions divided over bins by groups **(Figure 4b** and **inset)**. Despite analogous peaks in all groups, the spontaneous and 4-AP groups do not extend the right tail, as in the BiC and MK-801 groups, indicating recorded electrodes with low coherence values. On the other hand, the distributions of BiC and MK-801 groups are shifted to the right, indicating an increase of the spatial coherence for the average oscillatory activity in the whole OB neuronal circuit. These significant changes in the spatial coherence may set new standard values, especially in the GL layer (i.e., greater than 0.3), which meet the criteria for considering the GL as a morpho-functional basis of the spatial coding in the early processing of the sensory input^69^.

**Figure 4.**
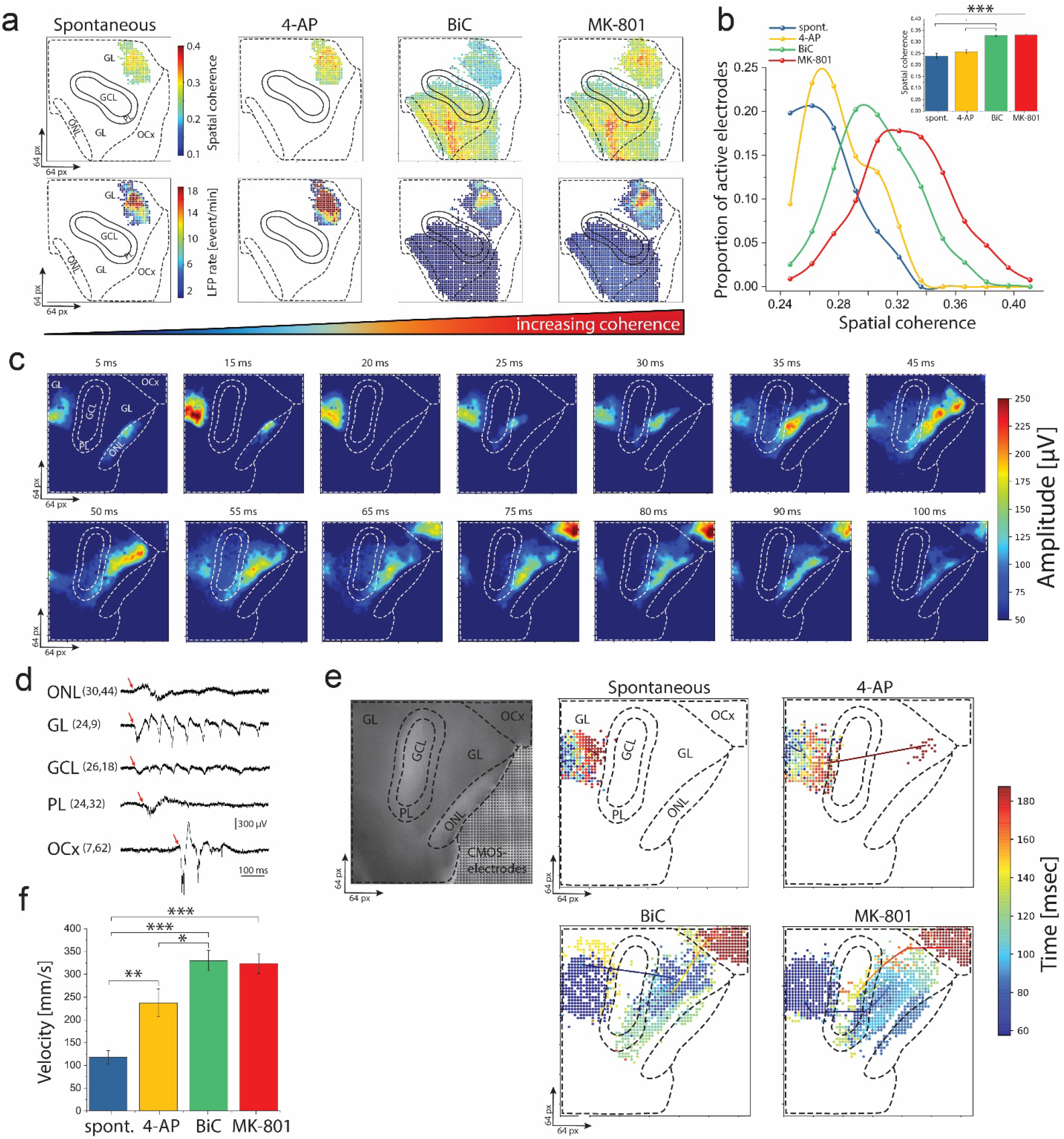
Topographical analysis of spatiotemporal dynamics of coding information. **a**) Spatial coherence maps displaying correlated pairs of recorded sites in the OB circuit. Maps on top showing the coherence, and at the bottom the maps of firing information encoded in field potentials. Spatial coherence increasing sequentially by pharmacological treatment of 4-AP, BiC, and MK-801, respectively, yielding more active electrodes encoding higher information of the firing rate of that spectral frequency component (i.e., in the GL). **b)** Spatial coherence quantification showing a significantly shifted distribution towards higher coherence values under drug-treatment compared with spontaneous activity and summarized in (**b**, inset) (\*\*\**p* < 0.001, ANOVA) (*n=24 slices from 8 mice*). **c**) Profile of propagation sequence of representative LFP event showing a defined initiation (ONL and GL) and dynamic propagation of the LFP firing patterns in the OB interconnected circuitry (sending output at the OCx). **d)** Temporal event traces corresponding to active recording sites in different OB layers. The red arrows point to the initiation time and shift to the right to indicate the propagation sequence (i.e., initiated in ONL-GL and terminates at OCx). **e)** Averaged CATs superimposed to the propagating active electrodes and the morphological structure of the OB layers. Time color-coded scale (*right*) indicates the CATs propagation profile and the firing electrodes of a single LFP event. **f**) Velocity of conduction of propagating LFP events is significantly increased when recording performed with 4-AP, BiC, and MK-801 compared to spontaneous (\**p* < 0.05, \*\**p* < 0.01, \*\*\**p* < 0.001 ANOVA) (*n=24 slices from 8 mice*).

In turn, this mapping may provide insights into a better understanding of large-scale circuit coding, decoding, and functional features of synchronized firing patterns in OB circuits.

### Localized initiation and large-scale propagation of synchronized oscillatory activity

In a multilayered OB circuit, the site of generation and spatial propagation of rhythmic electrical patterns are critical for the integrative activity of olfactory information processing. We used the simultaneous recordings from all OB layers to compute the initiation of the field potential signals and trace the spatiotemporal propagation of individual synchronized events. **Figure 4c** and **Supplementary Movie 2** show multi-frame pseudo-color images of the LFP oscillatory activity overlaid on the OB circuit’s corresponding anatomical regions. In the case of MK-801 treatment, the input signal is processed at the ONL. The oscillatory activity is initiated in the GL layer, propagates across GCL and PL layers to finally send output at the OCx layers. These characterized patterns sculpt complex propagation dynamics of the cellular mechanisms underlying the spatiotemporal coordination of information processing displayed by glomerular, mitral, and granule cells in the OB circuit.

Furthermore, **Figure 4d** shows the horizontal propagation of the LFPs across all OB layers. The dynamical propagation is illustrated as the time shift of the peak of the oscillatory events of exemplified signal traces selected from five electrodes located in different OB layers.

Next, to quantitatively investigate the propagation dynamics in detail and identify the ignition site of the location-weighted average of spatiotemporal LFP patterns, we employed a population parameter using the center-of-activity trajectories (CAT)^54^. We calculated the CAT mean for each electrode by integrating the total number of LFP events in 200 ms voltage-coded movies (see methods). We performed the CAT analysis on the spontaneous, 4-AP, BiC, and MK-801 groups that showed distinct ignition sites in the GL layer in all groups, with a dynamic population propagation towards the OCx in BiC and MK-801 (**Figure 4e**).

Furthermore, measuring a multi-site relation of spatiotemporal differences of occurring events allowed us to compute the instantaneous velocity of displacements of the CATs of LFP events in spontaneous and pharmacologically-induced groups to describe the dynamic of spreading activity within the interconnected layers of the OB circuitry. We found a range of propagation mean velocities of CATs ranged from (117 ± 17 to 323.67 ± 21.4 mm/s) in spontaneous and MK-801 groups, respectively (**Figure 4f**). These values are in the range of other previously reported for olfactory bulb velocity of conduction in rats (250-460 mm/s)^70,71^.

In sum, these results display the global dynamics of the OB population activity by incorporating the physical location of the electrode and the neuronal firing patterns of the multilayered OB circuitry. Also, illustrate evidence of spatial changes in interconnected neuronal populations of the bulbar rhythmogenesis upon blocking the dendrodendritic synaptic connections with MK-801. Similar results were also confirmed by previous in vivo reports on the role of NMDA receptor blockade for dendrodendritic inhibition concluded in smaller-scale electrophysiological and molecular readouts^72,73^.

### Multilayered OB functional decomposition for network-wide connectivity

The intricate spatiotemporal propagation patterns illustrated in the previous section involve diverse local neuronal connectivity necessary to synchronize the OB population. To unravel OB circuitry’s complexity to understand the underlying information processing, we have examined the emergence of inter-bulbar functional connectivity patterns as a network of pairwise firing electrode interactions. In this context, we computed the Pearson correlation between all LFP firing electrode pairs and performed the OB regional clustering to identify the level of synchronization (**Figure 5a**). Quantitatively, the cross-correlation increased significantly after pharmacological treatment with 4-AP, BiC, and MK-801 compared to the baseline spontaneous activity (**Supplementary figure 2**). Further, by tracing the emergence of functional links from spontaneous to MK-801-evoked activity, we found in the GL, predominant distinct populations of activated ensembles forming hub microcircuits and connecting OB subregions circuitry (**Supplementary figure 3**). This mesoscale rewiring of those new connections is signaled by the activity-dependent changes processed by the contemporary firing ensembles in the GL and determined the interareal coordination and reorganization of the population activity. These results illustrate individual neuronal ensembles’ contribution to the OB network synchrony in ever-available detail. Further, help to identify the critical OB subregions for optimizing network patterns in a large-scale spatial organization.

**Figure 5.**
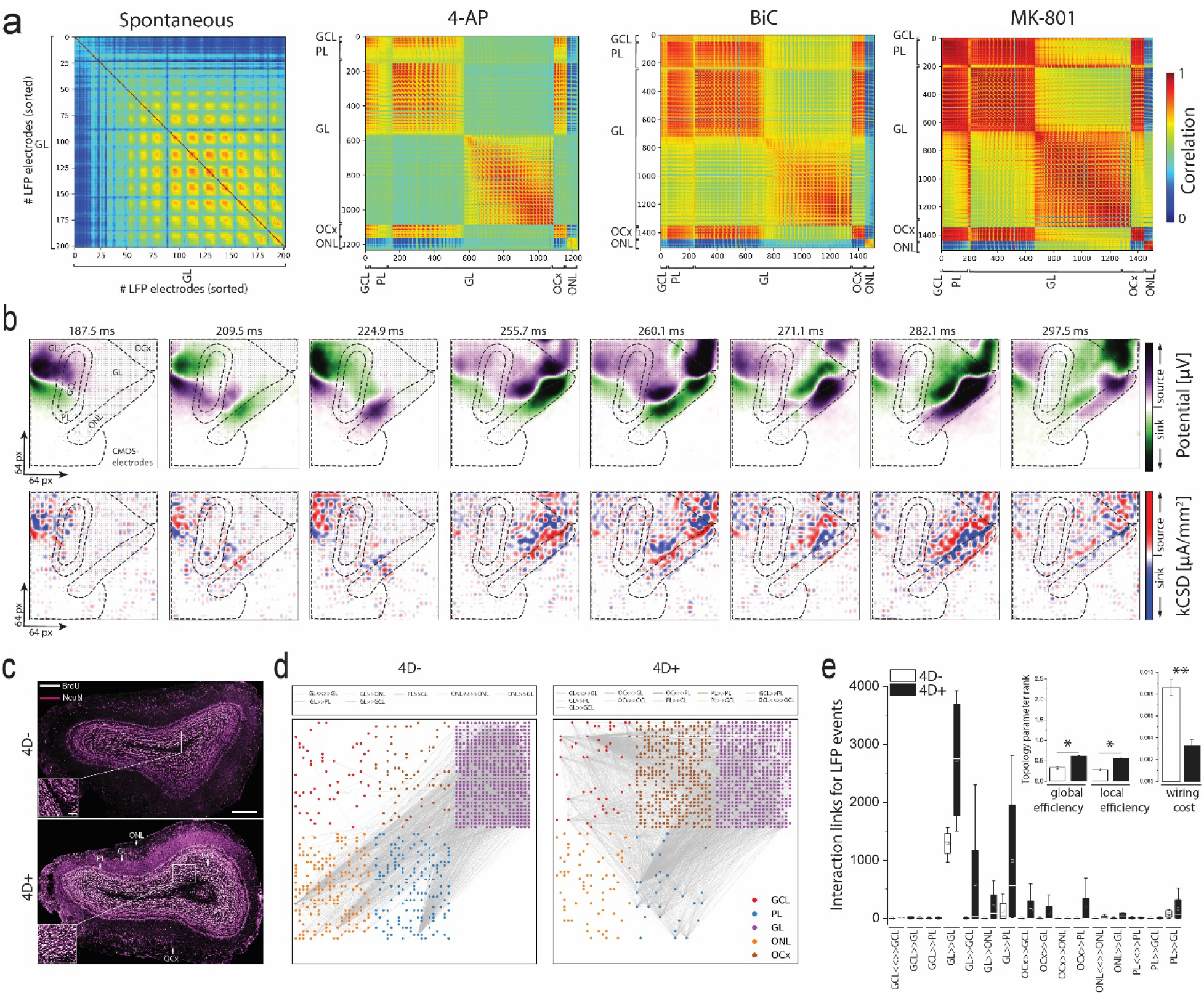
Large-scale functional connectivity analysis and validation on dynamical circuit remodeling. **a)** Functional connectivity matrices of different active regions of the OB circuit computed using the cross-correlation method for each pair of active electrodes in the 4096-array. In the spontaneous phase, only electrodes of the GL layer showed correlation to each other. The cross-correlation matrices in 4-AP, BiC, and MK-801 treatment indicate higher connectivity with more electrodes simultaneously fired and higher network synchronization than spontaneous recordings. **b)** The temporal evolution of a 2D kCSD construction (bottom) from the LFP recordings (top) for ongoing 100 ms activity. The kCSD indicates the sources (red tones) and sinks (blue tones) during the global field potential activation of the entire OB circuit. **c)** Evident OB neurogenesis effect in 4D+ mouse model as obtained from^45^. Confocal micrographs showing 4D+ slices with higher expression of NeuN+/BrdU+ of newborn integrated neurons mainly in the GCL compared to 4D-. Scale bars represent 500 μm (whole OB) and 100 μm insets. **d)** Direction-based functional connections of interactive pairs of firing electrodes in 4D− and 4D+ recordings computed using Granger causality and DTF algorithms. Grey links showing only 20% of the total detected links on 64×64 CMOS-recording array and are sorted in 6 clusters of the OB layers. **e)** Quantification of (D) showing significantly higher unidirectional and bidirectional interactive links in the 4D+ compared to 4D− (\**p* < 0.05, ANOVA). (E, inset) showing the effect of induced OB neurogenesis on the topological network parameters of efficiency (\**p* < 0.05, ANOVA) and cost of information processing in the entire OB circuit (\*\**p* < 0.01, ANOVA) (*n=12 slices from 4 mice*). 4D+ indicates higher network efficiency at a lower wiring cost compared to 4D− group.

### Identifying OB neuronal transmembrane current source and sink generators

Next, we further exploited our platform to pinpoint the high spatiotemporal resolution of local field generators (i.e., sources and sinks) in the inter-bulbar circuitry as emerged from the oscillatory activity of OB neuronal ensembles. Thus, we constructed bidimensional maps employing the kCSD method^58^ to estimate the average transmembrane currents extracted from the field activity recordings (**Figure 5b,** and **Supplementary Movie 3**). At the beginning of the LFP activation (*t=187.5 ms*), the KCSD maps (**Figure 5b, *bottom***) showed defined topographic activation features indicated by stronger and focused source generators (red color-coded in the GL and GCL layers with defined cellular identity (i.e., the microelectrodes underlying the transmembrane dipole). In contrast, the potential maps (**Figure 5b, *top***) showed general activation with no topographical differences in the GL and GCL layers. The computed propagating source-sink generators accompanied the oscillatory synchronized activation dynamically across OB layers. These current generators are eventually conveyed information at the OCx layers (*t=297 ms*) but provided more defined and sharper localization of the cellular neuronal activation (i.e., in the OCx, multiple sources revealed by kCSD while only sinks were detected in the potential maps). These results demonstrate remarkable advantages in defining and mapping disjoint sets of sinks and sources of the OB oscillatory activity compared to potential measures in terms of strength of neuroelectrical activation and spatial topographic patterns. This allows us to circumvent field potentials’ ambiguity and pinpoint the LFP generators at a high spatiotemporal resolution to capture with fidelity the correlated functional and individual cellular elements.

### Large-scale recordings validating enhanced OB-wide network connectivity and efficiency with adult neurogenesis

Next, to validate the significance of our large-scale recordings, we sought to examine the impact of ongoing neurogenesis on the dynamic of cross-scale OB circuitry and olfaction improvement. Thus, we used a genetically modified mouse model allowing the inducible expansion of neural stem cells by overexpression of Cdk4/cyclinD1 (4D+) in neural stem cells and using littermate (4D−) mice as negative controls^45^. Structurally, the 4D+ group has shown a neuronal expansion and increased neurogenesis indicated by the NeuN+/BrdU+ staining compared to the 4D− group (**Figure 5c**)^45^. On the other hand, using standard patch-clamp whole-cell recordings reported no significant functional and electrophysiological differences between 4D+ and 4D-group^45^. Thus, here, we obtained OB-wide patterns of activity from 4D+ and 4D-groups. Then, we applied the Granger causality as a connectivity method^74,75^ to quantify the directional information flow in the OB layers rendered by all pairwise electrode interactions (**Figure 5d**). The 4D+ group showed significantly higher unidirectional (GL→ GCL, OCx →GL, GCL, PL) and bidirectional (GL→GL, PL, ONL) interactions compared with the 4D-group (**Figure 5e**).

However, despite the localized functional induction by the new neurons in the GCL layer of the 4D+ group, the neurogenic effect significantly enhanced the global performance capacity of information processing in the entire interconnected OB circuitry, thus promoting network economy^76^. To this highlight, our results demonstrate the trade-off between network cost and efficiency indicated by the increased interconnection density between OB layers, reduced physical cost (i.e., wiring cost), and increased local and global efficiencies of information transfer of the spatially connected neuronal ensembles (**Figure 5e inset**) (see methods). Remarkably, our results also concluded network-wide functional remodeling properties that were unfeasible in smaller-scale electrophysiological methods.

## CONCLUSIONS AND OUTLOOK

The unmet need for large-scale multi-site bioelectrical imaging techniques limits the understanding of information processing in the olfactory system. It also hampers implementing olfactory biomimetic cell-based biosensors to exploit olfaction in fundamental applications. Dense microchips composed of thousands of electrodes could overcome this limitation and reveal the olfactory spatiotemporal dynamics. Thus, we have devised BIONICS (Biohybrid Olfactory Neural Circuit on a CMOS-chip), a novel platform with high-density CMOS-based active circuit architecture for simultaneously recording from thousands of OB neuronal ensembles to delineate large-scale spatiotemporal morpho-functional plasticity in the multilayered OB circuitry.

These recordings are facilitated by functionalizing the CMOS-chip surface with the adhesion-promoting molecule (PDLO). This allowed a homogenous and effective electrical coupling of the neuro-electrode-wide interface (i.e., generation of a high seal resistance) that fostered cellular activity stability and optimal SNR. Accelerating development in bioelectronics nose platforms hinges on the measurement technique and biosensing instruments as well as complex analytical tools for acquiring fundamental insight about the recorded biosignals. Thus, we employed rigorous algorithms and statistical analysis to catalyze significant parameters and insights from our multidimensional recorded data.

We have attributed the inhomogeneity and sparsity of OB cross-scale functional representations to their skewed lognormal LFP distributions driven by a balance between excitatory and inhibitory neuronal populations to provide network stability and reliable information processing for spatiotemporal coding.

We further characterized the firing patterns and their dynamical features across the OB-wide network by classifying the waveform shapes based on their rich inherent information in their nonsinusoidal oscillations. These results can inspire computational models and analytical methods to correlate these distinct waveforms to their underlying biophysical generators to describe disease conditions or behavioral states.

Also, facilitated by multi-site measurements of pharmacological-induced activity, we concluded the emergence of spatial network coherence, initiation of the LFP oscillations, and their cross-scale propagational dynamics at high spatiotemporal resolution.

The large number of recording sites in the CMOS-array confer sufficient access to simultaneous neuronal ensembles, enabling the construction of high-resolution 2D network-wide functional connectivity matrices and kCSD maps of local field potentials. Thus, providing a plausible method to probe the intrinsically sparse OB network connectivity and the contribution of local microcircuits and inhomogeneities in the LFP generation.

Our study suggests a new tool to address plasticity underlying olfactory spatiotemporal coding, not accessible with current imaging and electrophysiological techniques. Remarkably, we have specifically assessed the large-scale functional remodeling in the OB circuit that involved integrating newly generated neurons in the GCL layer of a mouse model with enhanced neurogenesis. We elucidated the induced functional neurogenic effect for enhancing OB network performance by integrating wide-effective information rendered by the topological organization and the local and global network efficiency. This would suggest a new generation of biologically inspired smart dynamical biomimetic systems and computational models capable of achieving faster and optimized olfactory information processing and enabling optimal selectivity and sensitivity for challenging applications.

In sum, our work has attempted to paint with a large-scale neurotechnological approach the breath and wonders of the chemosensory olfaction system. BIONICS has meticulously broken down the intricate firing patterns associated with olfactory code, promising progress, not only toward understanding neural processing of sensory information in the olfactory system but the biomimetic design of a bioelectronic nose and artificial chemosensory system development that may provide specific biomarkers for health and disease.

## Supporting information

Supplementary Movies description

## ACKNOWLEDGMENT

We would like to thank Prof. Dr. Gerd Kempermann, Dr. Caghan Kizil (DZNE, Dresden), and Dr. Alessandro Maccione (3Brain AG, Switzerland) for their insightful comments on the manuscript and the fruitful discussion. We also thank Dr. Daniel Wójcik and Mr. Władysław Średniawa (NENCKI Institute, Poland) for their support on the kCSD analysis. We are thankful to Dr. Sara Bragado Alonso for supporting experiments using the 4D line. We would also like to acknowledge the Animal platform of DZNE-Dresden (Dr. Alexander Garthe, Anne Karasinsky, and Sandra Günther) for their support. We also like to acknowledge Ms. Katarzyna Wiśniewska (Thames British School, Poland) and Ms. Brett Emery (DZNE-BIONICS lab, Germany) for proofreading and discussing the manuscript’s clarity and readability.

## AUTHOR CONTRIBUTIONS

HA: Project conceptualization, planning, performing & management, and writing the manuscript

FC: Provided the mouse line and data concerning the 4D overexpression system

DK: Performed OB slicing and technical support

SK: Performed part of experiments and analyzed the data

XH: Wrote the code for analysis and analyzed the data

All authors revised, reviewed, and approved the final version of the manuscript.

## Notes

### Competing Interest Statement

The authors have declared no competing interest.

