## Supplementary Figures for "Revealing Spatiotemporal Circuit Information of Olfactory Bulb in Large-scale Neural Recordings"

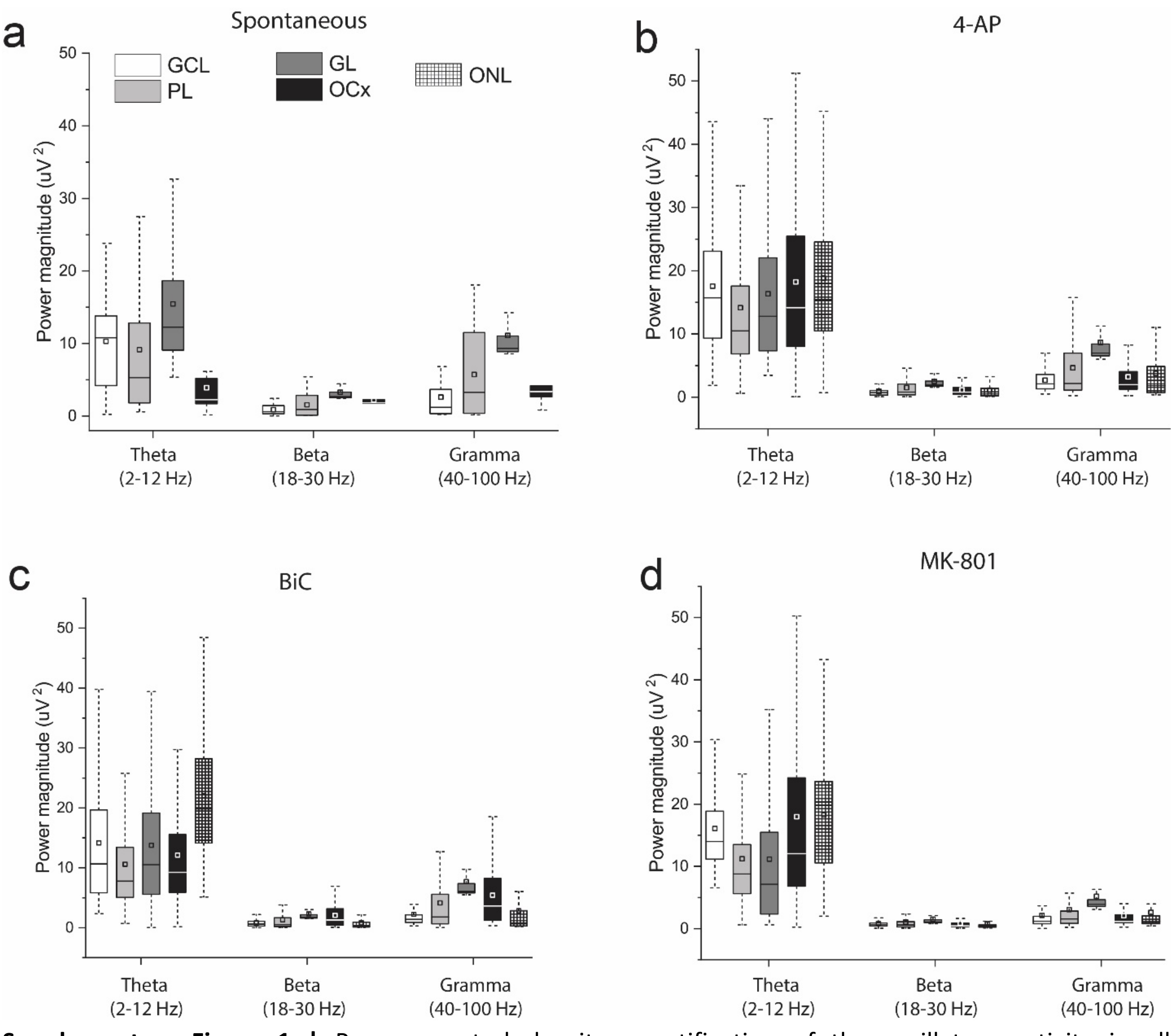


**Supplementary Figure 1 |** Power spectral density quantification of the oscillatory activity in all interconnected layers. The LFP signals' strength displayed in three major oscillatory Broadbands in **a)** spontaneous, **b)** 4-AP evoked activity, **c)** BiC-evoked activity, and **d)** MK-801-enhanced activity. *(n=24 slices from 8 mice)*.

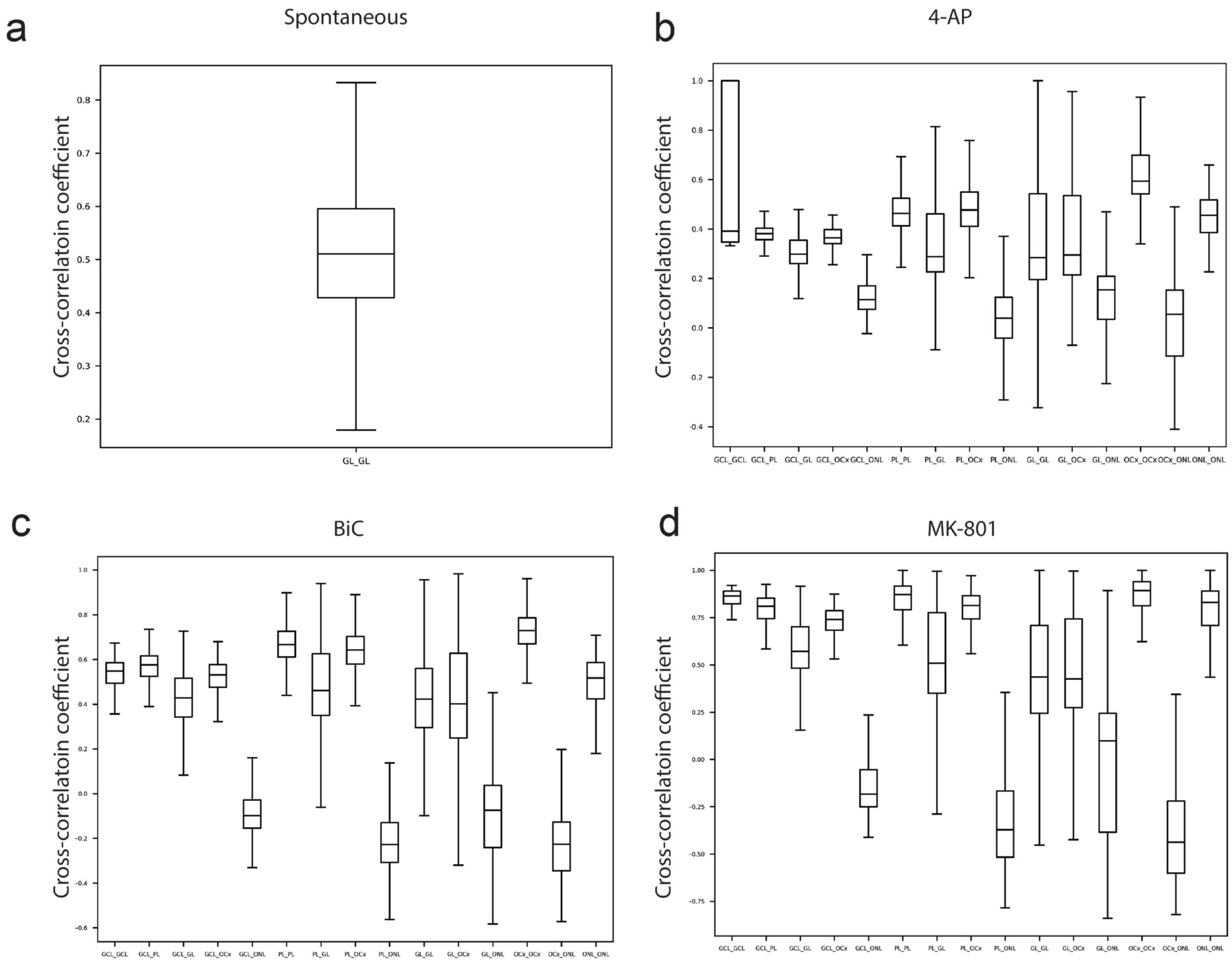


**Supplementary Figure 2|** Cross-correlation coefficient quantification. **a)** In spontaneous activity, only electrodes in the GL layer are firing LFPs and display correlation. **b)** 4-AP increased the firing discharges in the entire OB; thus, pairwise correlated electrodes indicated a higher correlation compared to spontaneous. **c)** Like the 4-AP phase, BiC induced a higher correlation of the firing electrodes, further strengthened under the MK-801 treatment shown in **d.** *(n=24 slices from 8 mice)*.

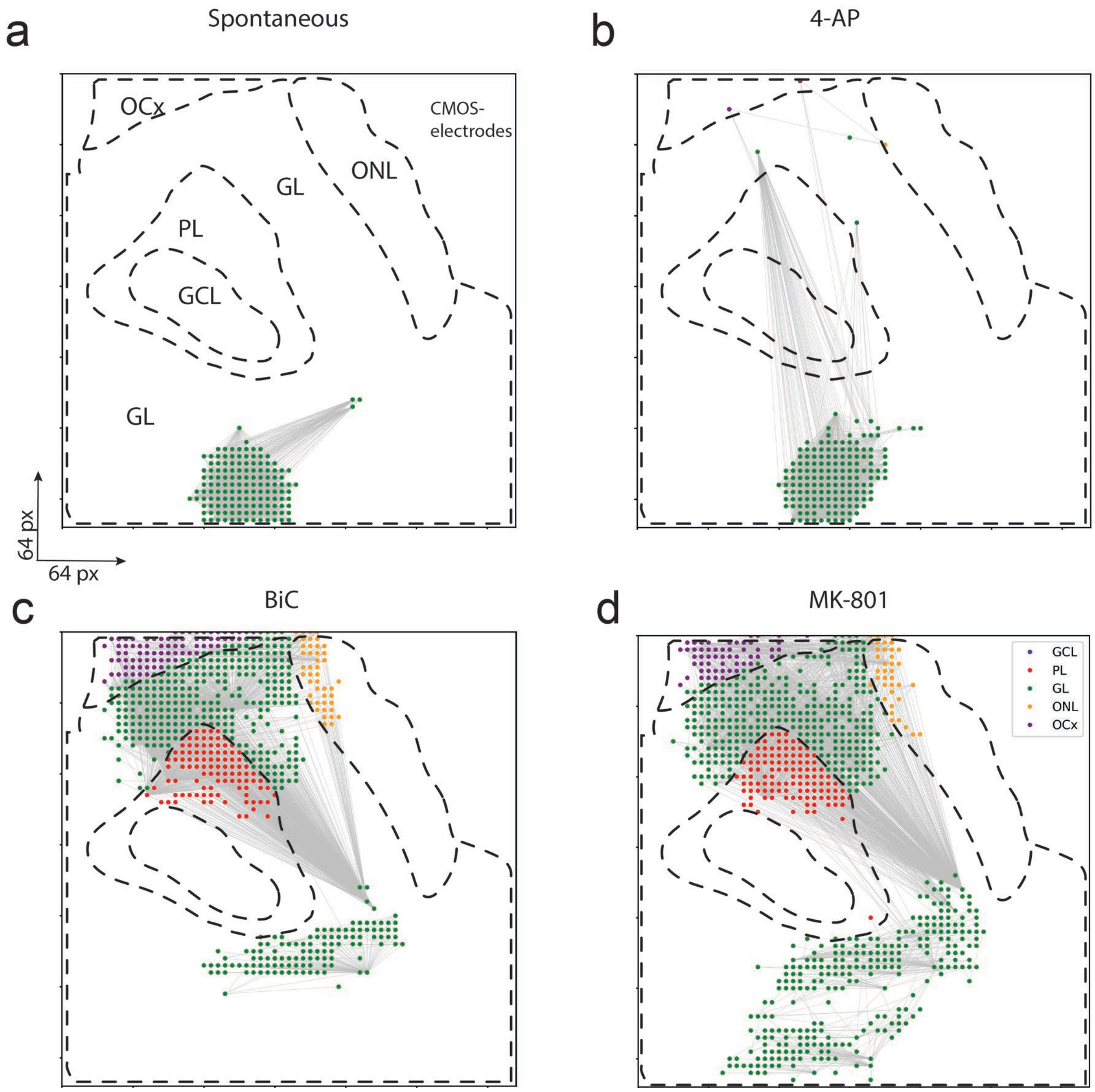


**Supplementary Figure 3|** Granger causality and network-wide remodeling with pharmacological treatment. The evolution of directed functional connections and network-wide rewiring computed with Granger causality and DTG algorithms. The links are emerging not only due to increasing firing electrodes but also their connection within the circuit remodeled based on the recording condition as in **a)** spontaneous, **b)** 4-AP, **c)** BiC and **d)** MK-801.
