## Supplementary Movies description for "Revealing Spatiotemporal Circuit Information of Olfactory Bulb in Large-scale Neural Recordings"

**Supplementary Movie 1:**

The movie clip displays real-time raw data (non-filtered, and no detected events 0.1-3 kHz), recorded with 14 kHz sampling frequency from full-frame 4096-array from the entire acute mouse olfactory bulb slice at 8 weeks old. The left video shows LFP oscillatory events across the entire array under the Mk-801 treatment. The right side displays five high signal-to-noise ratio selected active electrodes from different olfactory bulb circuitry layers. The shown event's real-time duration is 500 ms with a 20 ms bin size of the activity. The displayed video duration is 10 sec at frame rate = 25 fr/sec.

**Supplementary Movie 2:**

The movie clip displays a localized initiation and network-wide propagation of synchronized oscillatory events in the interconnected olfactory bulb slice. The shown spatiotemporal propagation overlaid on the slice indications renders the olfactory bulb structure underlying the firing event recorded from the entire CMOS-array (i.e., 64 x 64 electrodes). The shown event's real-time duration is 130 ms with a 5 ms bin size of the activity. The displayed video duration is 25 sec.

**Supplementary Movie 2:**

The movie clip displays bidimensional maps of transmembrane current source and sink generators in the entire active olfactory bulb circuitry. The top array shows the potential information of the oscillatory events. The bottom array displays the topographic features of the sources and sinks of the LFP activity computed with the kCSD method. The shown event’s real-time duration is 300 ms with a 20 ms bin size of the activity. The displayed video duration is 26 sec.
